# Model reduction permits Turing instability analysis of arbitrary reaction-diffusion models

**DOI:** 10.1101/213298

**Authors:** Stephen Smith, Neil Dalchau

## Abstract

Synthesising a genetic network which generates stable Turing patterns is one of the great challenges of synthetic biology, but a significant obstacle is the disconnect between the mathematical theory and the biological reality. Current mathematical understanding of patterning is typically restricted to systems of 2 or 3 chemical species, for which equations are tractable, but plausible genetic networks typically consist of dozens of interacting species. In this article, we suggest a method for reducing large biochemical systems to systems with 2 or 3 species which can then be studied analytically. We provide conditions to guarantee that the full system forms patterns if the reduced system does, and vice-versa. We confirm our technique with 3 examples: the Brusselator, an example proposed by Turing, and a biochemically plausible patterning system consisting of 17 species. These examples show that our method significantly simplifies the study of pattern formation in large systems.

## 1 Introduction

How cells co-ordinate with one another to form regular patterns of alternate differentiated states is a foundational question in developmental biology [1]. Establishing general rules that biochemistry can follow to enable pattern formation could impact on our ability to understand and cure developmental disorders [2], construct synthetic organs/organoids [3], or enable synthetic biology applications to utilise multicellular self-organisation [4, 5, 6]. While there are several mechanisms that are known to enable multicellular self-organisation of regular patterns, such as the *french flag* model [7], we focus here on *diffusion-driven instability* (DDI) first described by Alan Turing [8]. He proposed that two “morphogens” (intercellular signalling molecules) could enable tissues to produce regular patterns, and introduced a framework based on the reaction-diffusion equations that can establish when a given chemical system has pattern-forming potential. Later, Gierer and Meinhardt proposed that self-organisation requires a self-enhancing activator, which also up-regulates an inhibitor, forming a negative feedback, and further that the activator must diffuse more slowly than the inhibitor [9]. While an activator-inhibitor system is the simplest pattern-forming network, requiring only two chemical species but with differential diffusion, the introduction of a third (non-diffusing) species has been found to enable pattern formation when the morphogens have equal diffusion rates [10, 11].

Despite the theory of Turing patterns having existed since the 1950s, only much more recently has compelling evidence emerged that suggests that Turing patterns are responsible for pattern formation in natural biological systems, including digit patterning [12, 13] and fish skin colouring [14]. In most cases, it has been challenging to relate known biology involving many interacting species to simple 2- and 3-species networks for which analysis of DDI is more straightforward [13, 15]. As such, it remains the subject of debate as to whether the examples of biological pattern formation cited above actually depend on DDI, or might arise due to other reasons. To help understand the biochemical mechanisms that can result in biological pattern formation, several articles have proposed constructing synthetic biochemical networks that are engineered to specifically implement pattern-forming behaviours, some based on Turing instability [16, 17, 18, 19] but also other mechanisms [20, 21, 22, 23]. Libraries of biological parts/components have now been compiled that have been demonstrated to be functional in specific cellular systems that are frequently used in synthetic biology applications (e.g. *Escherichia coli, Saccharomyces cerevisiae).* Knowledge of the functioning of these components could then be utilised to demonstrate how manipulating kinetic parameters influence the conditions for DDI, and alter pattern wavelength, in predictable ways. Establishing a close relationship between theory and experiment would then provide further evidence that Turing’s mechanism can drive biological pattern formation. However, examples of synthetic biological circuits that can produce Turing patterns have yet to emerge, further raising the question of whether the Turing mechanism alone is sufficient to robustly generate regular patterns in a biological system.

Analysis of DDI for two species is now well-established [24], but quickly becomes more complicated with the introduction of additional (often non-diffusing) species [10]. While more theory and automated mathematical tools are now emerging that facilitate the analysis of DDI in general n-dimensional systems [25, 11], it still remains a challenge when the underlying system is nonlinear, as is typically the case in biological systems. Therefore, it is not uncommon to start with a more detailed mathematical description of a chemical system, then attempt to reduce it to a simpler form whilst retaining the majority of the behaviour of the detailed model [16]. However, little analysis has emerged that establishes whether the conditions of DDI are preserved during a model reduction, despite it being observed that model reduction can change the required diffusion ratio for pattern formation [10].

Many techniques have been established that reduce the size of ordinary differential equation (ODE) models, offering a starting point for interpreting the impact of model reduction on Turing pattern formation. Each technique is based on minimising the fidelity between detailed and reduced models with respect to a specific property (see [26, 27] for reviews of model reduction techniques). Some methods guarantee that equilibrium solutions (and their stability properties) are retained through a reduction, while others attempt to minimise the deviation of the transient behaviour of a specified model variable or variables, in response to a stimulus. Furthermore, some methods preserve the model co-ordinates/variables, while others do not. In biochemical systems, timescale separation techniques are often used, of which the most common are the quasi-steady state assumption (QSSA) and the quasi-equilibrium (QE) assumption [26]. Both involve removing species that are *fast,* substituting the concentration of these species for functions of the dynamic species that are derived from equilibrium relationships arising from the full system.

In this article, we investigate the question of whether model reduction can be applied to a chemical reaction network in a manner that preserves Turing pattern-forming behaviour. In section 2, we prove that if a reduced model forms patterns, then so does the corresponding full system (and vice-versa), given that some easily-checkable conditions are fulfilled. In section 3, we confirm our results on three separate CRNs, including the Brusselator, and a synthetic gene network with 17 species. These examples show that the method developed in this article allows for quick and easy Turing pattern analysis of aribtrarily large and complex chemical systems.

## 2 Theory

### 2.1 Background description of Turing instability

The majority of theoretical work on Turing patterns builds upon the classical reaction-diffusion equations for a chemical system undergoing diffusion. In the absence of convection/advection, the reaction diffusion equations are given by

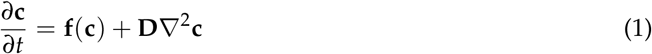

where 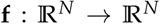 is in general a nonlinear system for the rate equations of a CRN involving *N* species (*X*_1_,…, *X*_*N*_), and **D** is a diagonal matrix containing the diffusion rates of each species. The ∇ operator describes the spatial derivatives in 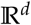, where *d* is the number of spatial dimensions. In 1d, this simply corresponds to 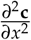

A Turing pattern arises when an equilibrium of the spatially homogoneous system (**c** = ĉ such that **f**(ĉ) = 0) goes unstable in the presence of diffusion. In our definition of a Turing pattern, this equilibrium is also assumed to be stable in the absence of diffusion. To analyse stability, we consider standard linear analysis of the system about the equilibrium *ĉ*. If 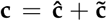 when 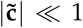, then (1) becomes

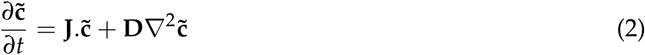

where **J** is the matrix of first-order partial derivatives of **f** with respect to each species *j*

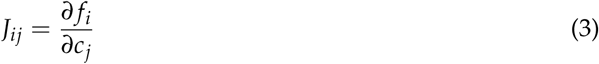

evaluated at **c** = **ĉ**.

To assess stability in the presence of diffusion, we consider how perturbations evolve over time. If *w*_*k*_(**x**) are the eigenmodes of the Laplacian operator ∇^2^, i.e. ∇^2^*w*_*k*_ = *η*_*k*_*w*_*k*_, then it has been shown that *η*_*k*_ ≤ 0 (with zero flux boundary conditions) [28]. Therefore, it is customary to let *η*_*k*_ = −*k*^2^, with *k* corresponding to the wavenumber of the eigenmode. As such, in 1d, on a domain *x* ∈ [0, *L*], there are solutions of the form

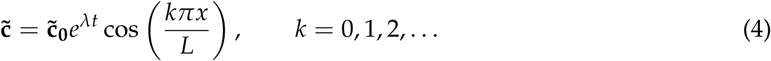

Accordingly, the original linearisation problem (2) translates into

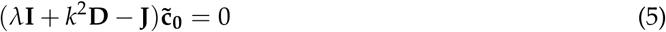

Therefore, we are interested in the eigenvalues of **J** − **D***k*^2^. If we denote by *λ(k)* the solutions of the eigenvalue problem associated with **J** − **D***k*^2^, then this gives rise to a dispersion relation

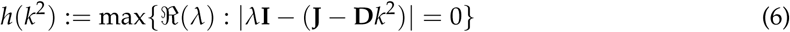

For Turing instability, we require that the system is stable in the absence of diffusion, which translates to eigenvalues at *k* = 0 all having negative real part. Additionally, we require the existence of at least one unstable wavenumber. i.e. there exists a wavenumber *k*\* such that there is a corresponding eigenvalue *λ*\* with positive real part.

### 2.2 Model reduction

We now consider a system of *N* species in which the first *M* species can diffuse with diffusion coefficients *D*_1_,…,*D*_*m*_, while the remaining species cannot. Accordingly, we describe a spatially inhomogeneous reaction-diffusion system as

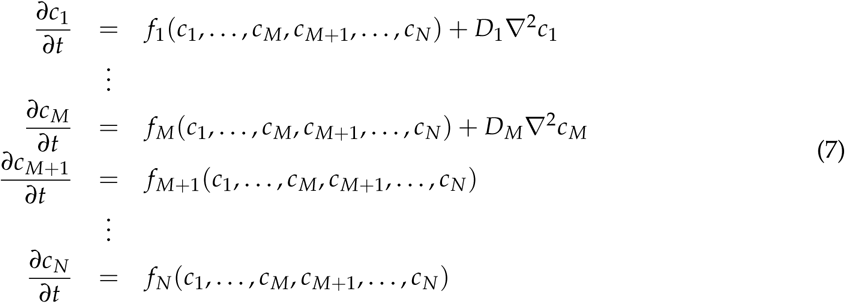

The associated spatially homogeneous system can therefore be written compactly as

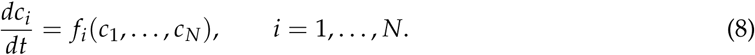

We further assume that there exists a non-negative spatially homogeneous equilibrium of (1) given by *ĉ*_*i*_ which satisfies:

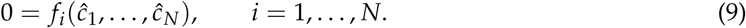

We now outline a strategy to reduce system (8) to a smaller system of *N*_*R*_ species, with *M* ≤ *N*_*R*_ < *N*, including all diffusible species. Without loss of generality, we assume that the reduced model consists of species *X*_1_,…,*X*_*N*_*R*__. The reduction is obtained by defining the functions 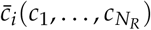, (*i* = *N*_*R*_+1,…,*N*), which satisfy:

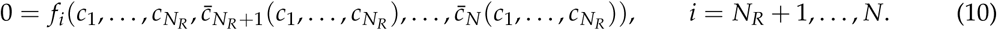

Intuitively, this amounts to solving the steady-state ODEs for the *removed* species, as functions of the *remaining* species, thereby eliminating *N* − *N*_*R*_ species from the system. Note that 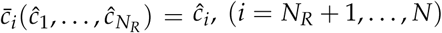. The reduced system of ODEs becomes:

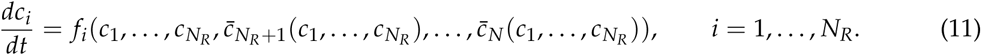

Using the chain rule, the Jacobian for system (11) is given by: evualated at <inline>, and the diffusion matrix is given by:

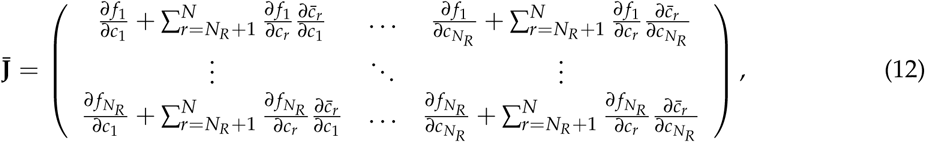

evualated at 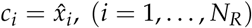 and the diffusion matrix is given by:

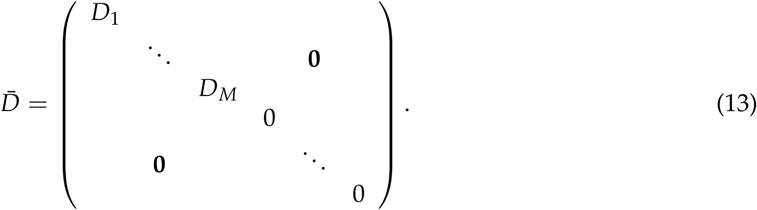

We now have two systems, (8) and (11), which are models of the same underlying process. To consider how Turing pattern formation is affected by the reduction from (8) to (11), we return to the mathematical conditions of pattern-forming behaviour introduced above.

We say a system is *pattern-forming* if there exist *k*_1_, *k*_2_,*k*_3_, *k*_4_ > 0 with *k*_1_ ≤ *k*_2_< *k*_3_≤ *k*_4_ such that all eigenvalues of **J** − *k*^2^**D** have negative real parts when *k* < *k*_1_ and *k* > *k*_4_, and there is a positive real eigenvalue when *k*_2_ < *k* < *k*_3_. This is a strict definition that explicitly excludes certain systems that are capable of forming patterns: (i) systems with patterns formed by Turing-Hopf bifurcations, (ii) systems that are unstable without diffusion, and (iii) systems that can form patterns on arbitrarily small length-scales (“noise-amplifying networks” [11]). Systems of type (i) are excluded because they can form either spatial patterns or temporal oscillations depending on the initial conditions, and so are not consistently pattern-forming; systems of type (ii) are excluded because they violate Turing’s concept of diffusion-driven instability; systems of type (iii) are excluded because they violate physical principles by permitting, for example, patterns on length-scales smaller than a molecule [10].

Knowing that we are interested in the behaviour of the matrix **J** − *k*^2^**D**, and its reduced counterpart 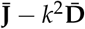, we note the following relationship between the full and reduced systems:

**Lemma 2.1.** |**J** − *k*^2^**D**| *and* 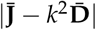 *have the same roots as functions of k*.

*Proof*. We define:

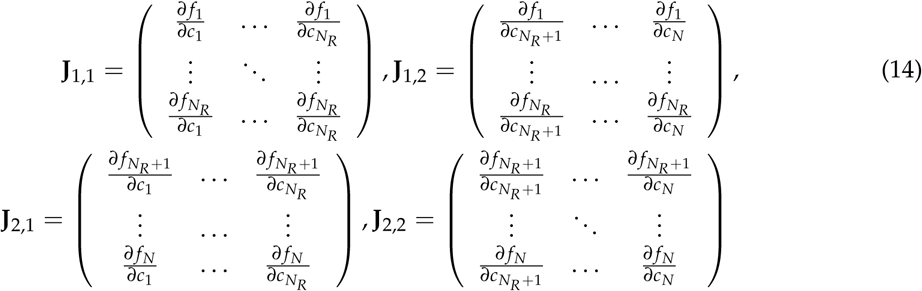

so that,

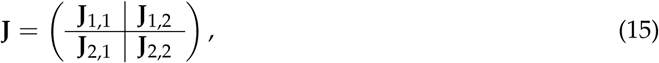

and we define:

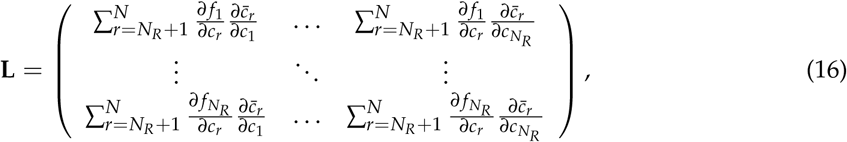

so that,

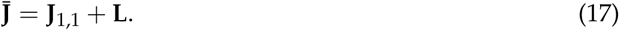

We now compare |**J** − *k*^2^**D**| and |**J** − *k*^2^**D**|. In the former case, we have that:

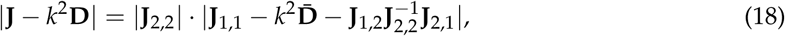

while in the latter case,

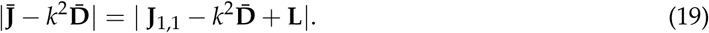

The two determinants are directly proportional if 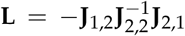. We observe that we can write 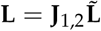, where,

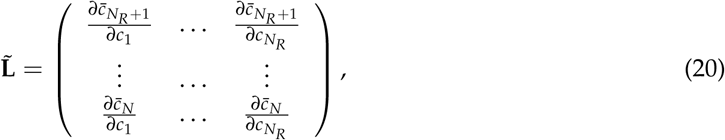

so that the condition for proportional determinants becomes that 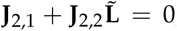, or algebraically, that

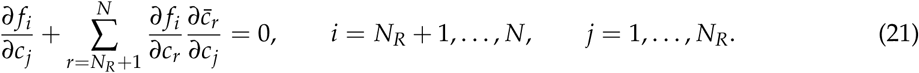

We note from Eq. (10) that, 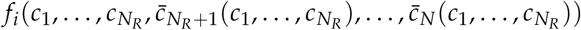, (*i* = *N*_*R*_ +1,&*N*) is a constant function of *c*_1_,…,*c*_*N*_*R*__, i.e.,

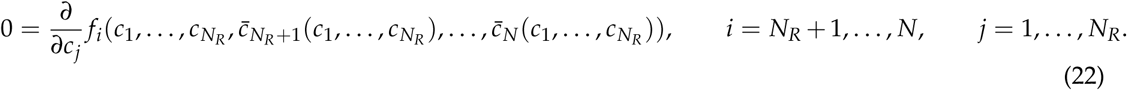

Expanding this gives precisely the condition (21). It follows that the determinants of the full and reduced systems are directly proportional, and consequently that any zero eigenvalues of the full system are replicated in the reduced system, and vice versa. □

In the case of two systems with two diffusible species, we can also make the following statement:

**Lemma 2.2.** *A system with two diffusible species (M = 2) is pattern-forming if it has the following properties*:

*(I) the system is linearly stable without diffusion (i.e. max(Re(eig(**J**))) < 0),*

*(II) the non-diffusible subsystem is either linearly stable without diffusion (i.e. max(Re(eig(**J**_2,2_))) < 0), or else non-existent (i.e. M=N=2)*,

*(III)* |**J** − *k*^2^**D**| *has two positive real roots (*k*_2_, *k*_3_ with 0 < *k*_2_ < *k*_3_) as a function of *k**.

*Proof.* By (I), all eigenvalues of **J** − *k*^2^**D** are negative when *k* = 0, and by continuity, also negative up to some *k*_1_> 0 with *k*_1_≤ *k*_2_. When *k* is very large, the characteristic polynomial of the system will have the form (*λ* − *k*^2^*D*_1_)(*λ* − *k*^2^*D*_2_)|**J**_2,2_ − *λ***I**| + *O*(*k*^2^) = 0 if *N* > 2, or else (*λ* − *k*^2^*D*_1_)(*λ* − *k*^2^*D*_2_) = 0 if *N* = 2. The eigenvalues of **J** − *k*^2^**D** will therefore converge to −*k*^2^*D*_1_, − *k*^2^*D*_2_ and (if *N* > 2) the eigenvalues of **J**_2,2_, which are all negative by (II). It follows that all eigenvalues of **J** − *k*^2^**D** are negative for sufficiently large *k* (say, larger than some *k*_4_≥ *k*_3_). Furthermore, since *M* = 2, |**J** − *k*^2^**D**| is a quadratic function of *k*^2^, it follows from (III) that |**J** − *k*^2^**D**| changes sign at *k*_2_ and *k*_3_, and nowhere else. Since there exist *k* < *k*_2_ and *k* > *k*_3_ both corresponding to all negative eigenvalues of **J** − *k*^2^**D**, it follows that there is at least one positive eigenvalue when *k*_2_ < *k* < *k*_3_. The system therefore satisfies all conditions required for pattern-forming behaviour.□

The combination of Lemmas 2.1 and 2.2 directly provides the conditions for which model reduction preserves pattern-forming behaviour. In particular, we have the following result:

**Lemma 2.3.** *If a full (reduced) system is pattern-forming, then the reduced (full) system is also pattern-forming if both the reduced (full) system and - if it exists - its non-diffusible subsystem is stable without diffusion*.

*Proof.* Conditions (I) and (II) of Lemma 2.2 hold by definition. Since the full (reduced) system is pattern-forming, there must exist distinct *k*_2_, *k*_3_> 0 such that 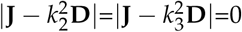. By Lemma 2.1, zeros of the full and reduced systems’ determinants coincide, so condition (III) of Lemma 2.2 also holds. Therefore the reduced (full) system is pattern-forming.

There are two important implications of this result for model reduction in practice. Firstly, if we reduce a large model and find a set of parameters for which the reduced model forms patterns, then we only have to check the Jacobian of the full model to find if it also forms patterns. This is useful because checking the stability of a Jacobian is computationally much simpler than finding largest real eigenvalues as functions of *k*, especially for systems with many species. Secondly, if the reduced model is stable for a region of parameter space, then the full model cannot form patterns in that region. This is useful because the stable region is typically large, and unstable regions frequently correspond to physically impossible parameter values, and so model reduction can be an efficient way of eliminating systems incapable of pattern formation.

In the next section we apply our technique to some example systems and confirm that our results hold.

## 3 Examples

### 3.1 Brusselator

One of the simplest chemical systems that is known to exhibit Turing patterns is the Brusselator, which in its original form is described by four reactions involving only two essential chemical species *X* and *Y* [29]

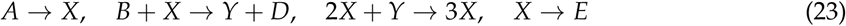

Here, the species *A, B, D* and *E* are explicitly included to ensure that mass is conserved. However, they are often removed during analysis as they do not contribute to the characterisation of the system behaviour, under the assumption that *A* and *B* are never depleted.

Following some debate over the chemical plausibility of reactions with more than two reactants, it was proposed in [30] that by introducing a third chemical species, the trimolecular reaction could be converted to a pair of bimolecular reactions

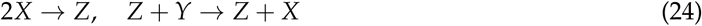

As the resulting bimolecular Brusselator system has not previously been analysed for Turing pattern formation explicitly, we applied our model reduction approach to determine conditions for which Turing instability is preserved. To simplify the reaction network whilst retaining full coverage of the space of possible behaviours of the bimolecular Brusselator system, we remove the non-essential species and remove two of the rate parameters, leaving:

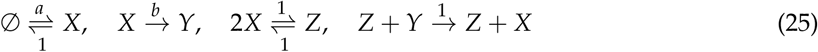

Assuming that X diffuses with unit rate, *Y* at a relative rate *D*_*Y*_ and *Z* is immobile, the concentrations of *X*, *Y* and *Z* for this system evolve as:

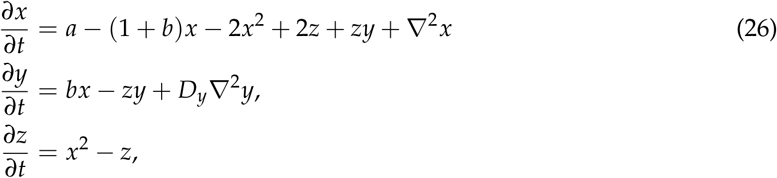

with equilibria 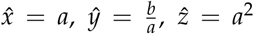. We perform a model reduction which removes *Z* from the system. As per our strategy, this is achieved by solving 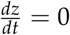 for 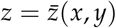. We get:

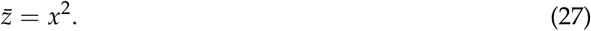

The reduced model is obtained by substituting Eq. (27) into Eq. (26):

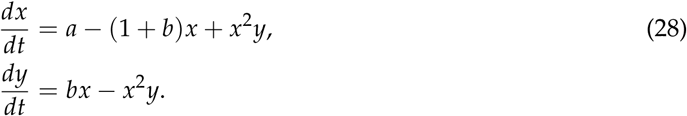

which recovers the reaction-diffusion equations for the classical Brusselator model.

In general, we find that parameter values which lead to patterns in the bimolecular Brusselator model also lead to patterns in the classical Brusselator model (Fig. 1 (a), (b)). To see this, we varied the parameter *b* and the diffusion constant *D_Y_*over large ranges, and compared the bifurcation diagrams. As we would expect from Lemma 2.3, these show that parameter values which lead to patterns in the reduced system also lead to patterns in the full system; correspondingly, parameters which lead to patterns in the full system lead either to patterns or instability in the reduced system. In Fig. 1 (c), (d) we show the patterns formed by the species X in systems (26) and (28) respectively.

**Figure 1:**
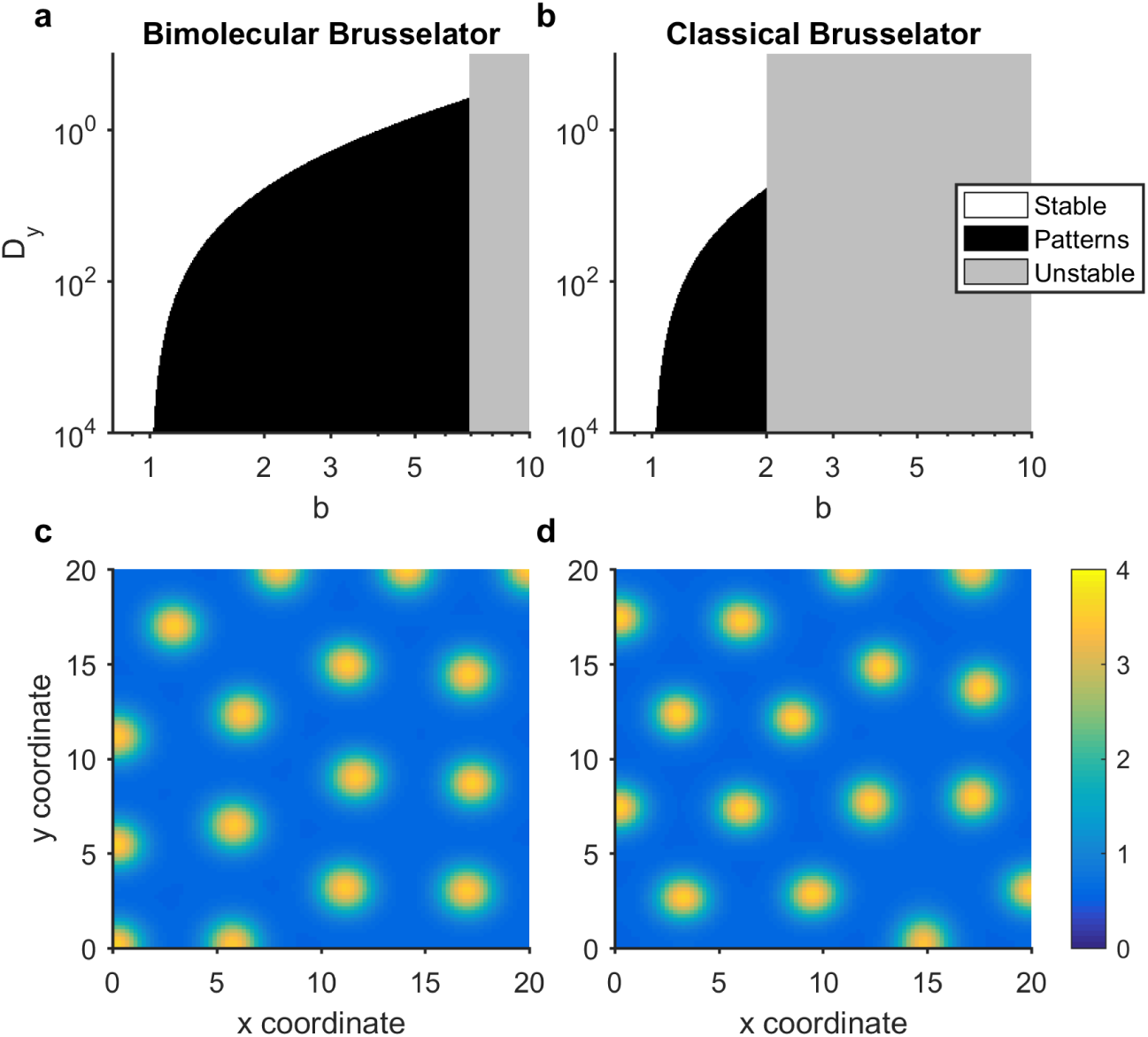
Model reduction of a bimolecular Brusselator mostly retains the bifurcation structure of the classical Brusselator model. Bifurcation diagrams are shown for (a) the bimolecular Brusselator (26) system and (b) the classical Brusselator (28) system. The clear area indicates parameter values of *b* and *D*_*Y*_ for which the homogeneous equilibrium is stable, while the black region indicates parameter values corresponding to Turing instability. The grey region indicates where the equilibrium solution is unstable in the spatially homogeneous scenario. (c),(d) Stable patterns formed by species X in the bimolecular Brusellator (26) and classical Brusselator (28) systems respectively. The parameter values used in these analyses were *a* = 1, *D*_*X*_ = 1, *b* = 1.88, *D*_*Y*_ = 10. Spatial simulations used a domain length of 20 (arbitrary units).

In Fig. 2 we show the dispersion relations for systems (26) and (28). As predicted by Lemma 2.1, while the relations themselves are different, they both change sign at the same values of *k*, implying that both systems will form patterns on the same wavelengths.

**Figure 2:**
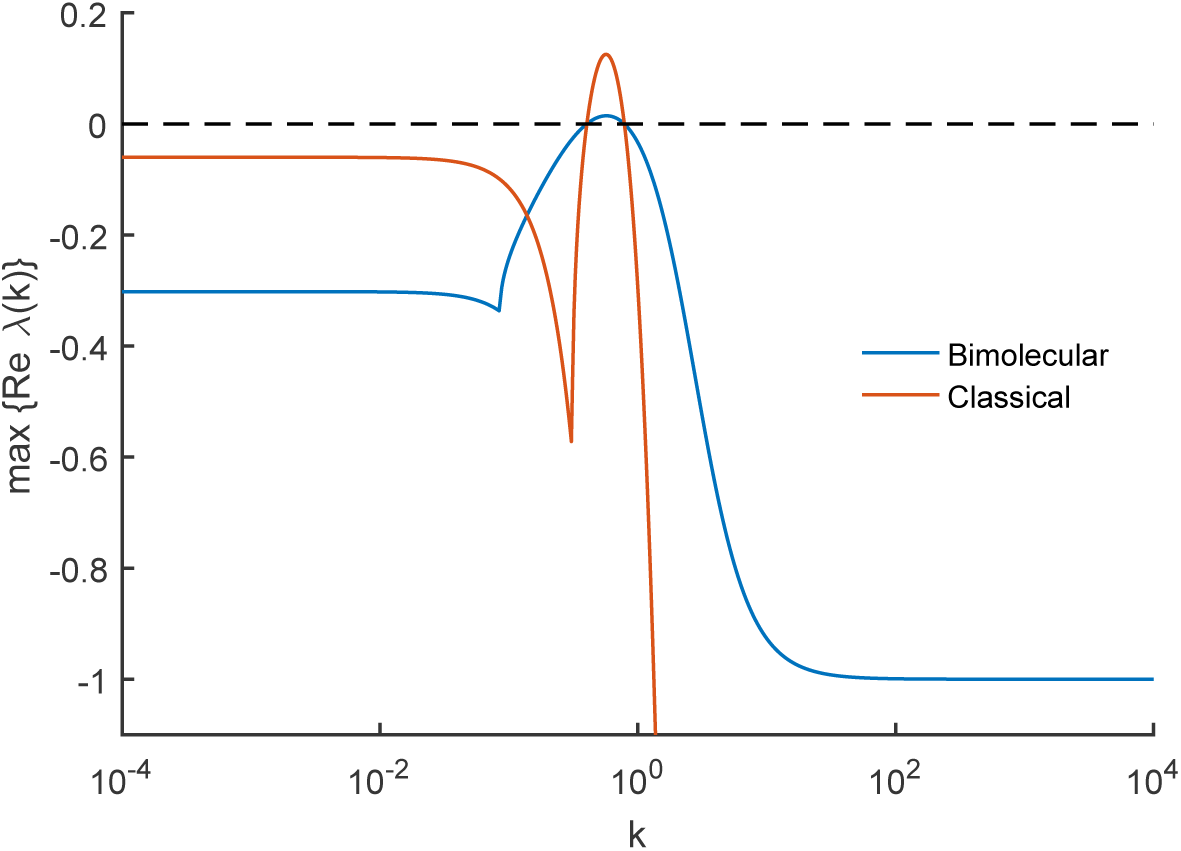
Dispersion relations for the bimolecular and classical Brusselators. Dispersion relations are shown for the bimolecular Brusselator (26) system and the classical Brusselator (28) system. Although the two plots are different in general, they both cross the zero line at the same points. Parameter values are as in Fig. 1.

### 3.2 Turing’s example

We next considered a larger example that is closely related to one proposed by Turing [8]. It consists of species *X, Y, W, C*, and *C*′, and concentrations *x, y, w, c,* and *c′* respectively. *X* and *Y* can diffuse with diffusion coefficients *D*_*X*_ and *D*_*Y*_, and the reactions are given by:

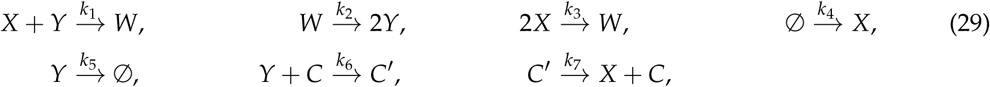

We note that the *C* and *C′* are related via a conservation law, and so we substitute *c′* = *c*_Tot_ − *c* (*c*_Tot_ constant), which leads to four independent ODEs that completely characterise the deterministic behaviour of the system:

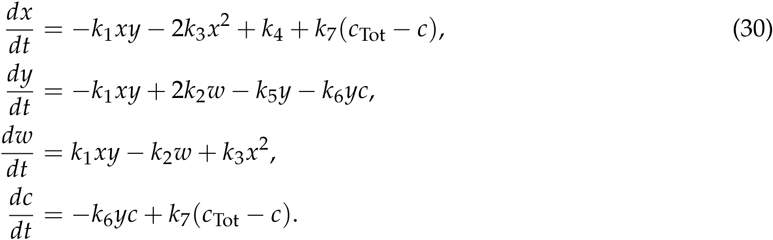

To demonstrate how the model reduction can be applied to different extents, we reduce this system to both 3- and 2-species system approximations. First, we eliminate *c* by solving 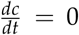 for 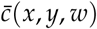, which gives

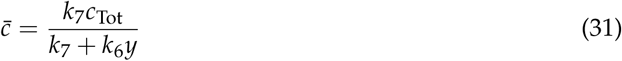

Substituting 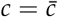; in system (30), we obtain a 3 species system defined by

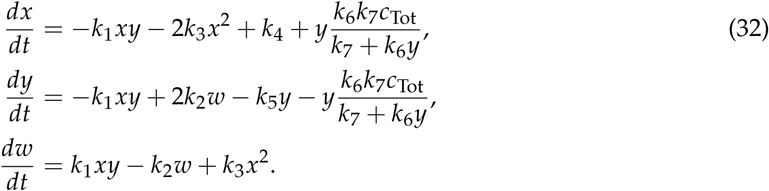

Next, we eliminate *w* by solving 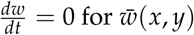, which gives

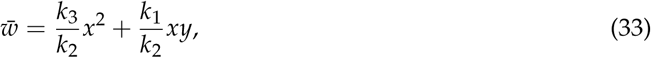

Substituting 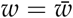 in (32), we obtain a 2 species system defined by

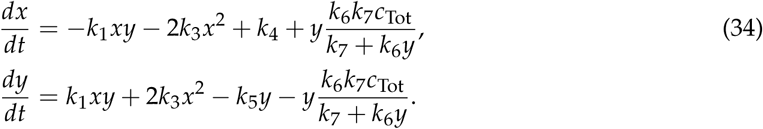

We therefore arrive at three models of system (29) with varying levels of dynamical complexity. A *complete* model is described by 4 species, whereas two successive equilibrium assumptions applied to C and then *W* generate two simpler models. To demonstrate the equivalence of Turing instability (Lemma 2.3) across these models, we illustrate bifurcation diagrams of the full system (Figure 3 (a)), and the reduced 3 (Figure 3 (b)) and 2 species (Figure 3 (c)) systems. These show that the 4 and 3 species models have indistinguishable parameter-dependent behaviour, while the 2 species model is unstable for a region of parameter space where the other models are stable. We note that this unstable region prevents the 2 species model from forming patterns when the diffusion rates of *X* and *Y* are equal (*D*_*X*_ = *D*_*Y*_ = 1), though such equal diffusion rates can produce patterns for the 3 and 4 species models. We also observe that pattern-forming parameters in the 2 species system also lead to patterns in the larger systems (Figure 3 (d)-(f)). This is similar to the situation observed for the Brusselator, whereby model reduction leads to a shrinkage of the parameter space that produces patterns. However, we have not established whether this is generally the case following equilibrium-based model reduction.

**Figure 3.**
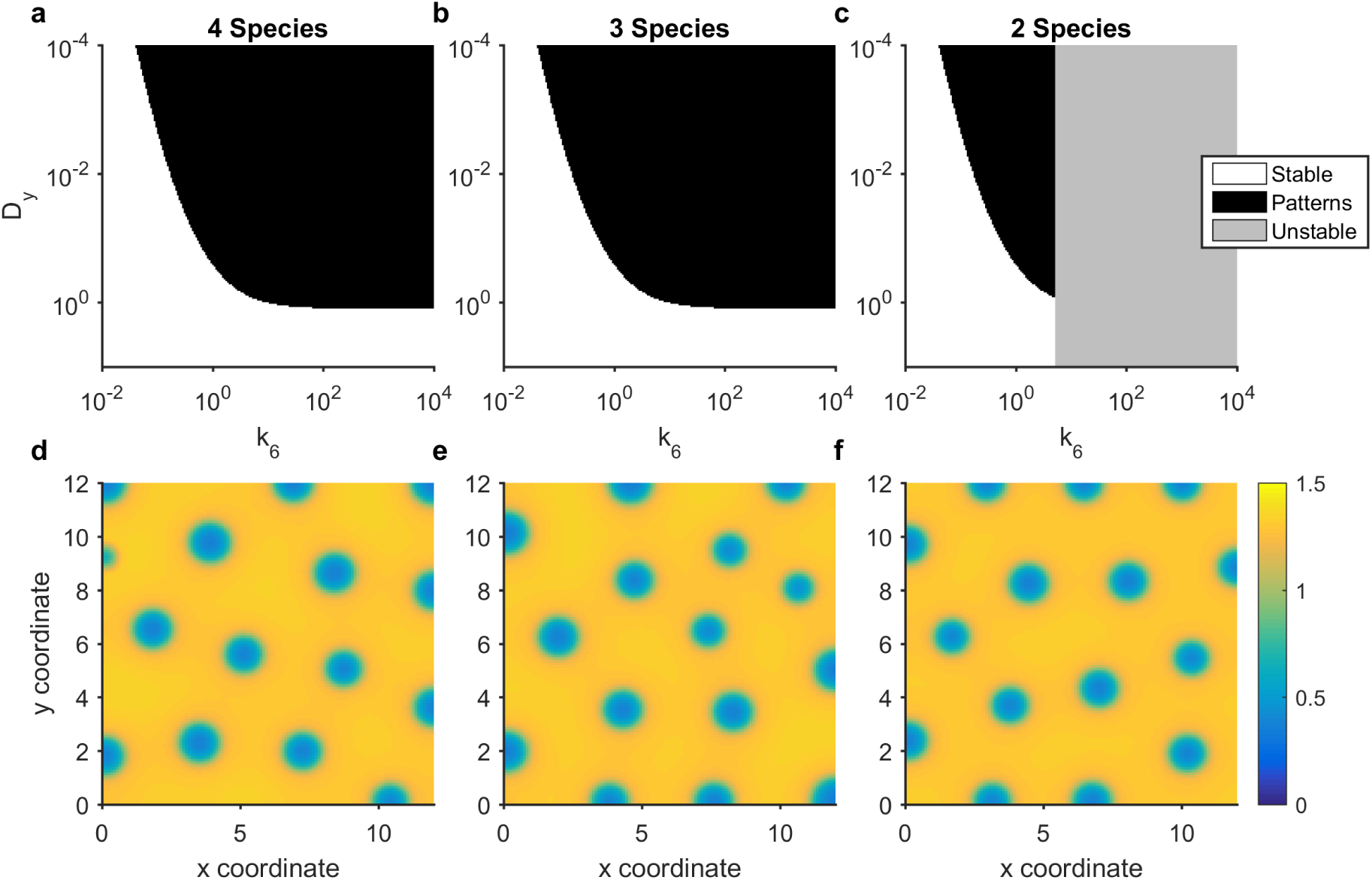
Turing pattern analysis for a reaction network from Turing [8]. (a-c) Bifurcation diagrams for 4- (30), 3- (32) and 2-species (34) systems. In all cases, the parameter values used were *k*_1_ = 2, *k*_2_ = 0.2, *k*_3_ = 0.01, *k*_4_ = 0.08, k_5_ = 0.04, k_7_ = 2, *c*_Tot_ = 6, *D*_*X*_ = 1. (d-f) Stable patterns formed by species X in the 4- (30), 3- (32) and 2-species (34) systems respectively. In all cases, the parameter values used were as in (a-c) but additionally *k*_6_ = 3.37, *D*_*Y*_ =0.04, domain length= 6.

In Fig. 4 we show the dispersion relations for the 4- (30), 3- (32) and 2-species (34) systems. As predicted by Lemma 2.1, while the relations themselves are generally different (although the 3- and 4- species relations are near-indistinguishable), they all change sign at the same values of *k*, implying that all systems will produce patterns on the same lengthscales.

**Figure 4.**
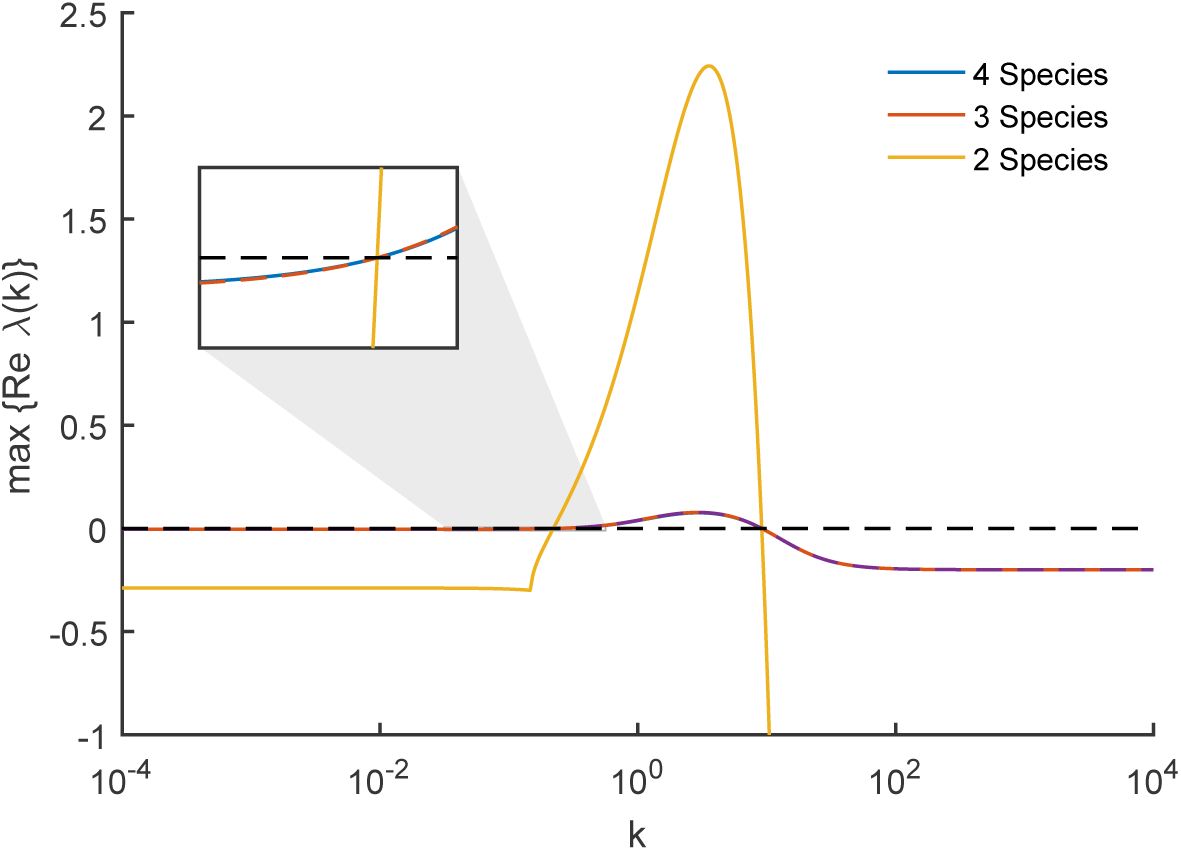
Dispersion relations for the reaction network from Turing [8]. Dispersion relations are shown 4- (30), 3- (32) and 2-species (34) systems (blue, red and yellow respectively). Although the plots are different in general, all three cross the zero line at the same points. An inset plot shows the crossing of the zero line at the lower value of *k*. Parameter values are as in Fig. 3.

### 3.3 A synthetic gene circuit

In our final example, we consider a much larger system consisting of 17 species, which is based on the synthetic gene circuit proposed in [18]. While this publication presents only a theoretical analysis of the synthetic gene circuit, it represents a biologically plausible approach to realising a synthetic cellular Turing patterning circuit in live cells. The synthetic gene circuit is arranged in an activator-inhibitor network, whereby the intercellular signalling molecule Acyl Homoserine Lactonase (AHL) plays the role of a short-range activator, and hydrogen peroxide gas (H_2_O_2_) plays the role of a long-range inhibitor. Activation is achieved by AHL binding a constitutively expressed LuxR receiver protein, forming an activating complex for PLux promoters, which are placed upstream of coding sequences for the AHL synthase luxI [31] and the H_2_O_2_-producing ndh. The inhibitory loop is formed by an H_2_O_2_-sensitive topA promoter stimulating production of the AHL lactonase aiiA, which degrades AHL [32], thus inhibiting its action.

In [18], it is shown that a model containing 5 variables (but analysis over 4 variables due to the presence of a conservation law) can give rise to diffusion-driven instability for certain parameter choices. Already, analysis of Turing instability is made challenging by virtue of there being more than 2 essential dependent variables. One might categorise their model as having intermediate complexity, as a simpler model could be arrived at by considering only the concentrations of the diffusive signals AHL and H_2_O_2_. In contrast, a more complex model might be considered that describes more of the intracellular components, and complexes between them, directly.

Here, we show that the bifurcation properties of models of the synthetic gene circuit in [18] are preserved across models of varying complexity. To demonstrate this, we start by considering a model described by elementary chemical reactions, as follows:

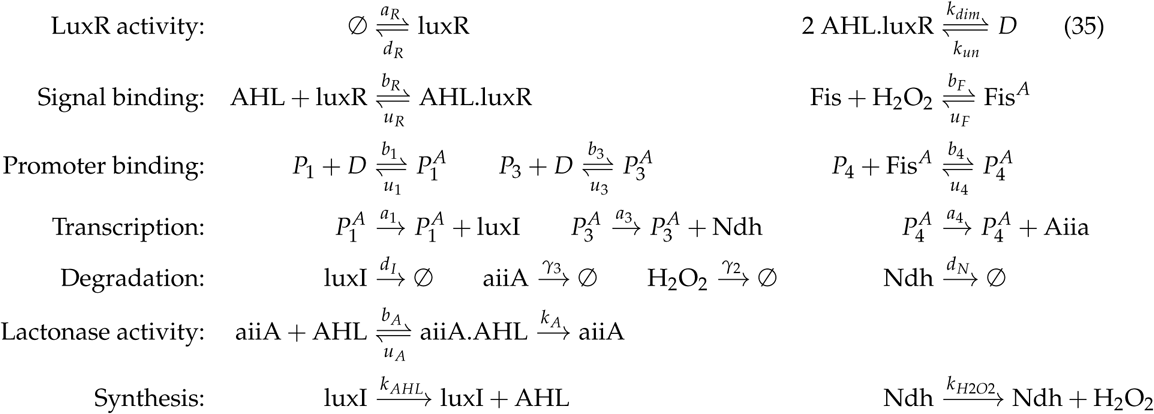

For brevity, we do not write out the full system of reaction rate equations here (although code is available from the authors upon request). We reduce the full system of equations to one of intermediate complexity consisting only of AHL, H_2_O_2_, AHL.luxR, and aiiA (as considered in [18], whose concentrations we write as *L*, *H, P* and A respectively. The reduced ODEs are:

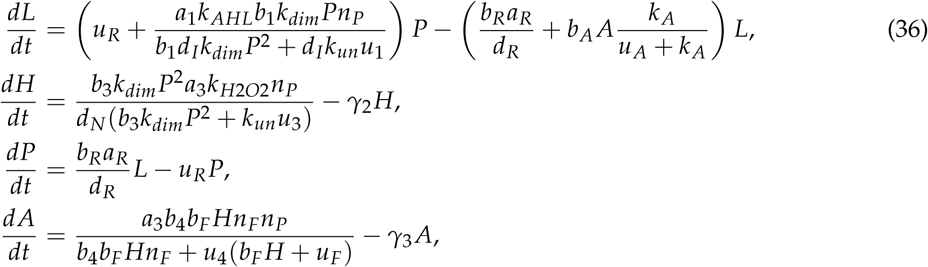

where *n*_*p*_ is the total promoter concentration and *n*_*F*_ is the total *Fis* concentration. While this system is very similar to the four species model studied in [18], there are some minor differences, but we nevertheless still find that Turing instabilities arise.

Finally, we can further reduce the system of intermediate complexity to a system comprising only the diffusive molecules AHL and H_2_O_2_:

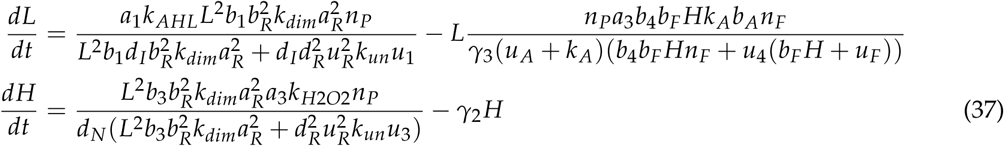

In Fig. 5 (a)-(c), we show bifurcation diagrams of the full system (35), the four species model (36) and the two species model (37). In this case, we find that the diagrams for the full (35) and intermediate (36) complexity systems are identical, while the diagram for fully reduced system (37) shows an unstable region of parameter space where the larger models are stable (in accordance with Lemma 2.3). All three models have identical pattern forming regions. In Fig. 5 (d)-(f) we show stable two-dimensional patterns of [AHL] in each system, which illustrates how patterns of a similar wavelength emerge. This is confirmed by the dispersion relations shown in Fig. 6, which show that each system’s dispersion relation changes sign at precisely the same wavenumbers (in accordance with Lemma 2.1).

**Figure 5.**
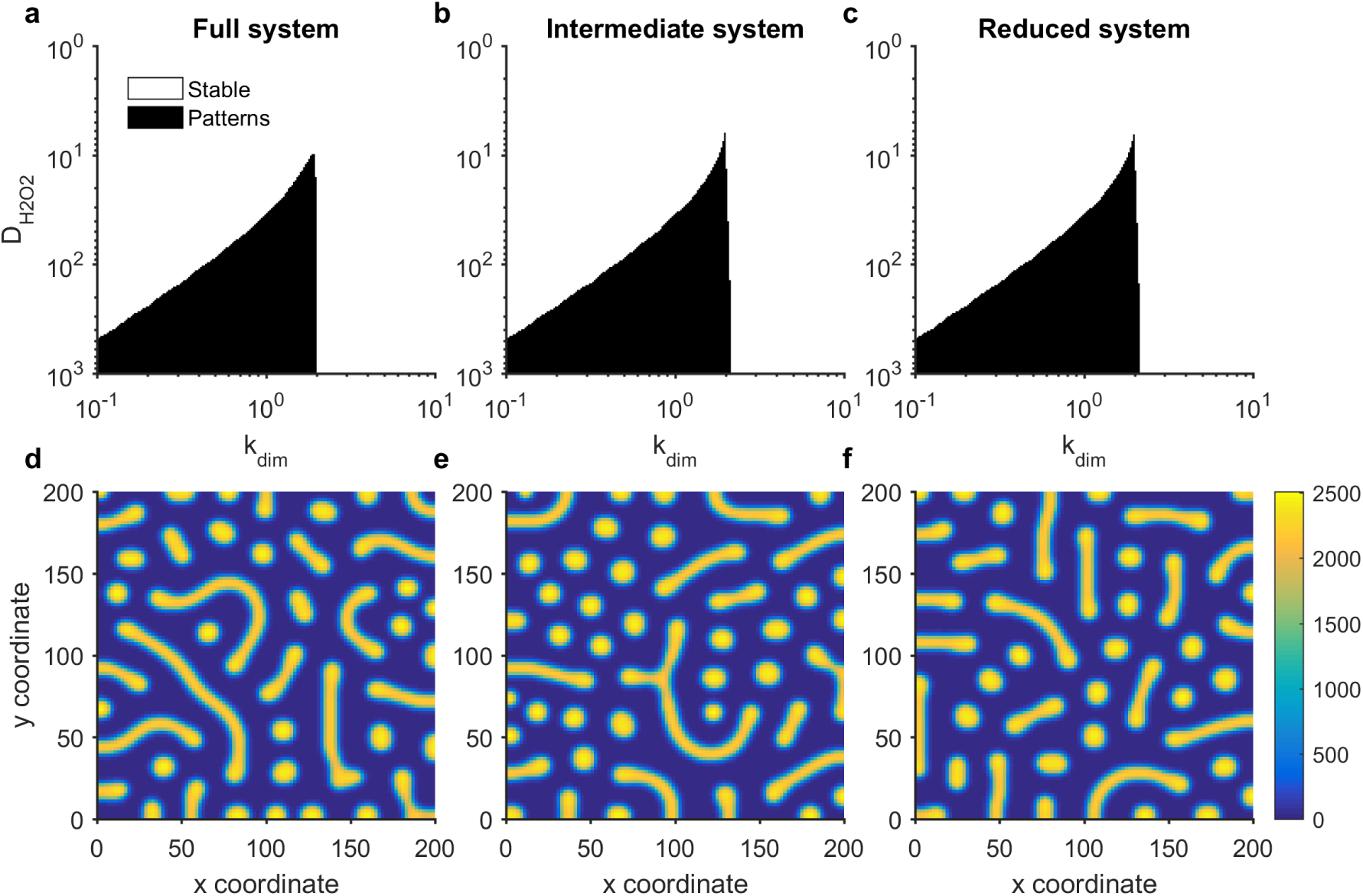
Turing patterns are robust to reductions of a model of a synthetic gene circuit with intercellular signalling. (a-c) Bifurcation diagrams for systems (35), (36) and (37) respectively. (d-f) Stable patterns formed by species AHL in systems (35), (36) and (37) respectively. Parameter values: *a*_1_ = 2142, *a*_3_ = 1190, *k*_*AHL*_ = 2, *k*_*H2O2*_ = 0.057, *b*_1_ = 0.156, *b*_3_ = 0.03, *b*_4_ = 0.25, *b*_*F*_ = 2, *d*_*N*_ = 2, *d*_*I*_ = 2, *d*_*R*_ = 2, *γ*_2_ = 2, *γ*_3_ = 2, *b*_*R*_ = 0.0156, *u*_*R*_ = 2, *u*_*A*_ = 2, *k*_*A*_ = 2, *b*_*A*_ = 0.0117, *a*_*R*_ = 0.5, *k*_*un*_ = 2, *n*_*F*_ = 2, *n*_*P*_ = 2, *u*_1_ = 2, *u*_3_ = 2, *u*_4_ = 2, *u*_*F*_ = 2, *D*_*AHL*_ = 1; in panels d-f, *k*_*dim*_ = 2, *D*_*H202*_ = 100, domain length= 100.

**Figure 6.**
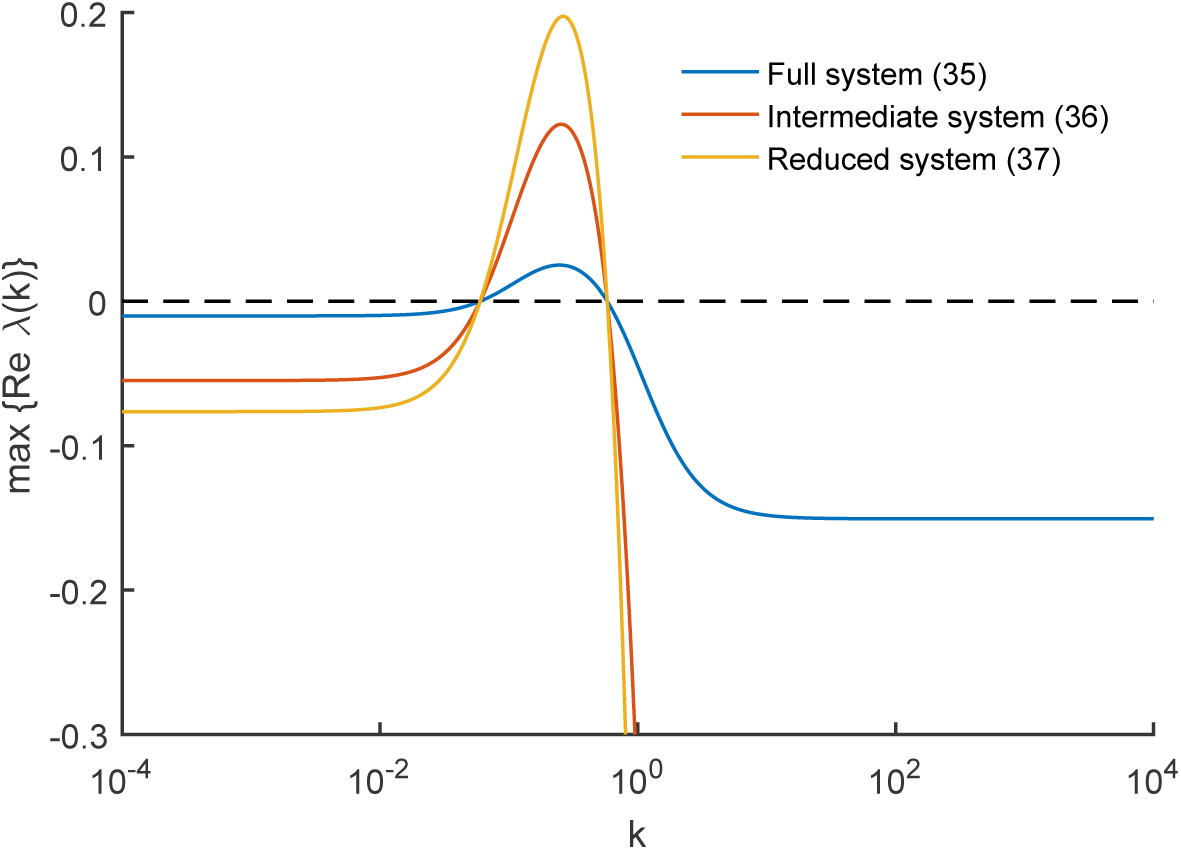
Dispersion relations for the synthetic gene circuit model. Dispersion relations are shown for systems (35), (36) and (37) (blue, red and yellow respectively). Although the plots are different in general, all three cross the zero line at the same points. Parameter values are as in Fig. 5.

## 4 Discussion

In this article we have proposed a technique for reducing a large chemical system to a small one, in a manner that preserves pattern-forming behaviour as far as possible. In essence, the reduction relies on a quasi-steady-state approximation (QSSA), since it assumes that the concentrations of the removed species can be written in terms of the remaining species without reference to time. However, the comparison to the QSSA is slightly disingenuous, since all species (not just the removed subset) must be in steady-state for stable Turing patterns to form, and the QSSA is normally applied to time-dependent systems. Comparison with other model reduction techniques is also difficult, since unlike typical approaches, our strategy is not necessarily interested in preserving the correct dynamical behaviour of the full system: indeed, we make no claims that the reduced model is an accurate description of the original system. The reduced model is derived with one aim in mind: to help find parameters of the full model which can lead to Turing patterns, or to help prove that none exist.

In that respect, our technique is very successful. We have shown that, if we can find a pattern-forming parameter set for the reduced system, then it is simply a matter of checking the stability of the Jacobian of the full system to determine whether it, too, forms paterns with those parameters. Furthermore, if we can find a region of parameter space for which the reduced system is stable, then we know for certain that the full system cannot form patterns in that region.

The power of our technique is demonstrated very well on system (35): a biologically plausible system consisting of 17 species and 31 reactions. At face-value, it is impossible to know whether this system is capable of forming patterns, and, if so, which parameters correspond to pattern-forming behaviour. By performing a dramatic reduction from 17 to 2 species, we quickly found regions of parameter space corresponding to pattern-forming and stability in the reduced model. Our results prove that these regions necessarily correspond to potential-pattern-forming and no-pattern-forming respectively in the full system. The fact that both systems generate near-indistinguishable patterns is an added bonus. Temporal dynamics are not conserved, which can be seen in the larger eigenvalues of reduced systems (Figures 2, 4 and 6), which is known to correlate with faster pattern emergence [33]. However, this is not surprising. Our model reduction technique is to simply assume that certain species equilibrate infinitely fast, and so the overall dynamics of reduced systems will be faster in general.

Overall, our results provide a quick and rigorous way to check for pattern-forming behaviour in large biochemical networks. While we do not attempt to automate this process here, such automation could be of serious utility to synthetic biologists in their attempts to find and synthesise genetic networks capable of forming stable patterns. The results in this article will also be of general interest to those in the reaction-diffusion field, since they provide a means to extend the current analytical tools developed for 2 or 3 species Turing-patterning systems to systems composed of an arbitrary number of species.

